# Sequential MAVS- and MyD88/TRIF-signaling triggers anti-viral responses of tick-borne encephalitis virus-infected murine astrocytes

**DOI:** 10.1101/2020.06.30.177485

**Authors:** Luca Ghita, Veronika Breitkopf, Felix Mulenge, Andreas Pavlou, Olivia Luise Gern, Verónica Durán, Chittappen Kandiyil Prajeeth, Moritz Kohls, Klaus Jung, Martin Stangel, Imke Steffen, Ulrich Kalinke

## Abstract

Tick-borne encephalitis virus (TBEV), a member of the *Flaviviridae* family, is typically transmitted upon tick bite and can cause meningitis and encephalitis in humans. In TBEV infected mice, *mitochondrial antiviral signaling protein* (MAVS), the downstream adaptor of *retinoic acid inducible gene I-like receptor* (RLR)-signaling, is needed to induce early type I interferon (IFN) responses and to confer protection. To identify the brain resident cell subset that produces protective IFN-β in TBEV infected mice, we isolated neurons, astrocytes and microglia and exposed these cells to TBEV *in vitro*. Under such conditions, neurons showed the highest percentage of infected cells, whereas astrocytes and microglia were infected to a lesser extent. In the supernatant (SN) of infected neurons, IFN-β was not detectable, while infected astrocytes showed very high and microglia low IFN-β production. Transcriptome analyses of astrocytes implied that MAVS-signaling was needed early after TBEV infection. Accordingly, MAVS-deficient astrocytes showed enhanced TBEV infection and significantly reduced early IFN-β responses. At later time points, moderate amounts of IFN-β were detected in the SN of infected MAVS-deficient astrocytes. Transcriptome analyses indicated that MAVS-deficiency negatively affected the induction of early anti-viral responses, which resulted in significantly increased TBEV replication. Treatment with MyD88 and TRIF inhibiting peptides reduced late IFN-β responses of TBEV infected WT astrocytes and entirely blocked IFN-β responses of infected MAVS-deficient astrocytes. Thus, upon TBEV exposure of brain-resident cells, astrocytes are important IFN-β producers that show biphasic IFN-β induction that initially depends on MAVS- and later on MyD88/TRIF-signaling.

## Introduction

Tick-borne encephalitis virus (TBEV) is a neurotropic positive-strand RNA virus belonging to the *Flaviviridae* family. This virus is closely related to other Flaviviruses such as West Nile virus (WNV), Japanese encephalitis virus (JEV), dengue virus (DENV) and others (1). TBEV is a constantly spreading zoonotic pathogen in Europe and Asia. The virus is transmitted via tick bites or consumption of infected milk (2, 3). Between 2012 and 2016, a total of 12,500 cases of TBEV infection were reported in Europe (4) and more recently this number increased up to 15,000 per year (5). Mostly, TBEV infection is associated with moderate disease, whereas in 10% of the cases a severe disease course is observed that culminates in the development of neurological symptoms such as encephalitis, meningitis and paralysis (6–9). The mortality rate ranges between 0.5 and 30%, depending on the virus strain. 30 to 60% of patients who recovered from TBEV induced encephalitis develop neurological sequelae (6–9). The increasing numbers of cases of TBEV induced encephalitis and the expansion of TBEV-affected areas in Europe enhance the medical relevance of TBEV. A vaccine is available, however, it is often not recommended outside of known endemic areas (10). Treatment options are not causative and limited to supportive care. This is due to the limited understanding of TBEV pathogenesis and immunity (11).

Upon bites by TBEV infected ticks, the virus is able to reach the central nervous system (CNS), whereas the detailed mechanism of infection is not fully understood. Currently it is believed that locally infected immune cells such as myeloid cells and lymphocytes migrate into the CNS by crossing the blood brain barrier, thus allowing the virus to access the CNS (12, 13). Within the CNS, TBEV productively infects neurons and astrocytes, which leads to the development of neuropathology (12). CNS-resident cells are able to sense virus RNA through *pattern recognition receptors* (PRRs) and then mount anti-viral responses (14). The PRRs mainly involved in the sensing of RNA viruses include the endosomal *Toll-like receptors* (TLR) and the cytoplasmic *retinoid acid-inducible gene-I* (RIG-I)-*like receptors* (RLR) (14–16). While TLRs are differentially expressed on CNS resident cell subsets, RLRs were shown to be ubiquitously expressed (17–19). Upon sensing of virus RNA within the cytosol, RLRs including RIG-I, LGP2 and MDA5 trigger downstream signaling through the adaptor molecule *mitochondrial antiviral signaling protein* (MAVS) (20). Triggering of MAVS induces activation of the transcription factors IRF3 and NF-κB, which leads to the induction of protective type I interferon (IFN) responses and expression of anti-viral genes (21).

Studies on TBEV pathogenesis revealed a crucial role of RLR signaling in the initiation of anti-viral immune responses. Upon infection with TBEV or Langat virus (LGTV), a naturally attenuated relative of TBEV, MAVS deficient mice (MAVS ko) showed enhanced susceptibility to lethal infection (22). Moreover, it was shown that LGTV infected MAVS ko animals presented with enhanced viral loads in the CNS. The lack of the MAVS adaptor molecule was correlated with an increased LGTV replication within neurons and astrocytes (22). Recently, it became evident that astrocytes actively participate in generating protective immunity upon TBEV infection. Cytokines and chemokines were shown to be released from astrocytes, thus contributing to the induction of an anti-viral state (23). Nonetheless, the astrocytic intrinsic mechanisms that initiate anti-viral responses upon TBEV infection are not well understood. Here we showed that neurons, astrocytes and microglia are productively infected upon *in vitro* exposure to TBEV, whereas astrocytes are the main producers of IFN-β protein. Transcriptome analyses revealed that the mechanisms underlying astrocytic responses to TBEV were strictly dependent on MAVS-signaling at early stages of infection, as MAVS ko astrocytes showed increased infection and reduced IFN-β production. Nevertheless, at later stages of infection, MAVS ko astrocytes mounted moderate IFN-β responses indicating the contribution of MAVS-independent mechanisms, which turned out to be MyD88/TRIF-signaling.

## Results

### Among brain-resident cell subsets, astrocytes are major IFN-β producers upon *in vitro* TBEV infection

To investigate the permissiveness of murine CNS-resident cells to TBEV infection, primary murine neurons, microglia and astrocytes isolated from C57BL/6 (WT) mice were exposed to TBEV Neudoerfl strain in cell culture at MOI 1. After 12 hours of incubation, all three cell types showed TBEV infection, as indicated by replication foci that were detected in perinuclear regions of the cytoplasm (Fig. 1a). Approximately 40% of the neurons displayed such replication foci, whereas amongst astrocytes and microglia only 26% and 18% of the cells were infected, respectively (Fig. 1b). Quantification of IFN-β protein levels in the SNs revealed that at 12 hours post infection (hpi) astrocytes produced significantly higher amounts of IFN-β than neurons and microglia (Fig. 1c). Thus, upon TBEV exposure, neurons showed a high percentage of infected cells and produced very little amounts of IFN-β, whereas astrocytes showed lower percentages of infected cells and produced the highest amounts of IFN-β of the analysed cell subsets.

**Figure 1:**
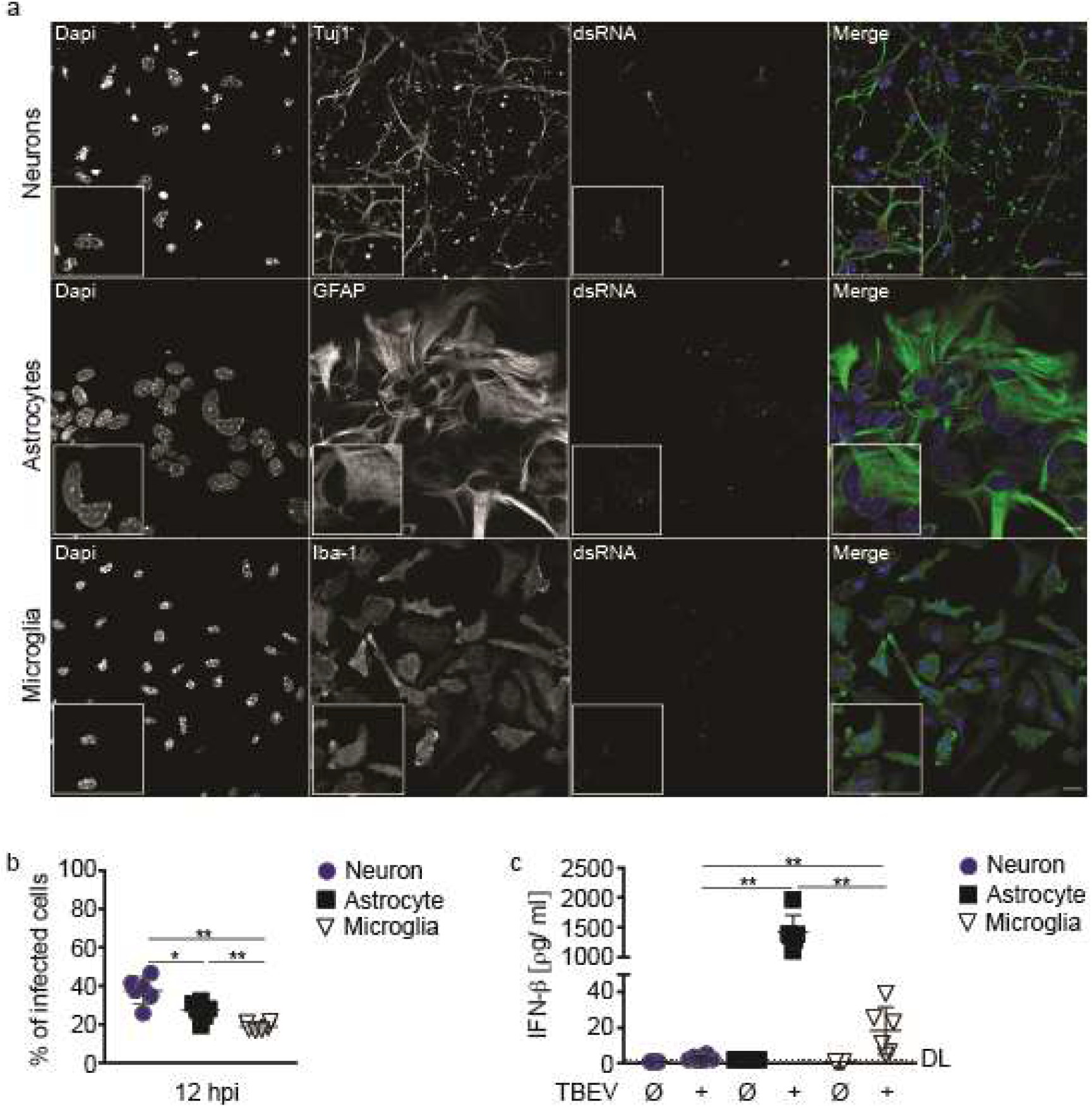
CNS-resident cell subsets show diverse responses to *in vitro* TBEV infection. Primary murine neurons, astrocytes and microglia isolated from C57BL/6 (WT) mice were infected with TBEV Neudoerfl strain at MOI 1. **a**, At 12 hpi cells were immunolabelled with either anti-Tuj1 (neurons), anti-GFAP (astrocytes), or anti-Iba1 (microglia) and anti-dsRNA (J2, Scicons English and Scientific Consulting, TBEV replication foci) and counterstained with DAPI. Lower-left corner shows enlarged picture of region of interest (white square). Scale bar 20 μm. **b**, Quantification of TBEV infected neurons, astrocytes and microglia at 12 hpi displayed as percentage of infected cells (n=6, combined data from 3 independent experiments). **c**, IFN-β protein levels in the SN of cultured primary neurons, astrocytes and microglia at 12 hours post TBEV infection was determined by ELISA (n≥ 5, combined data from 3 independent experiments). DL= Detection limit (0.94 pg/ml), error bars indicate mean ± SD; *P< 0.05, **P< 0.01; Two-tailed Mann-Whitney test.

### MAVS-signaling drives early innate responses in TBEV infected astrocytes

Transcriptome analyses of TBEV infected primary astrocytes verified the astrocytic identity of the analyzed cells (Supplementary Fig. 2a), whereas 12 hpi 458 differentially regulated transcripts were identified, of which 439 were up- and 19 downregulated when compared to mock treated cells (Fig. 2a). The upregulated transcripts encoded mostly interferons as well as interferon stimulated genes, while the downregulated transcripts were none immunoregulatory genes. A gene enrichment analysis of the 458 regulated genes indicated that TBEV infected astrocytes strongly upregulated genes involved in IFN-signaling pathways, including IFN genes as well as genes involved in the RLR-signaling pathway (Fig. 2b). Furthermore, normalized gene counts for *Ddx58, Eif2ak2*, and *Ifih1* confirmed the induction of RLR-signaling related genes (Fig. 2c,d,e). Collectively, these data indicated that astrocytes respond to TBEV infection by upregulation of IFN-dependent antiviral genes, and that RLR-signaling might be of key relevance for virus sensing and the induction of anti-viral responses.

**Figure 2:**
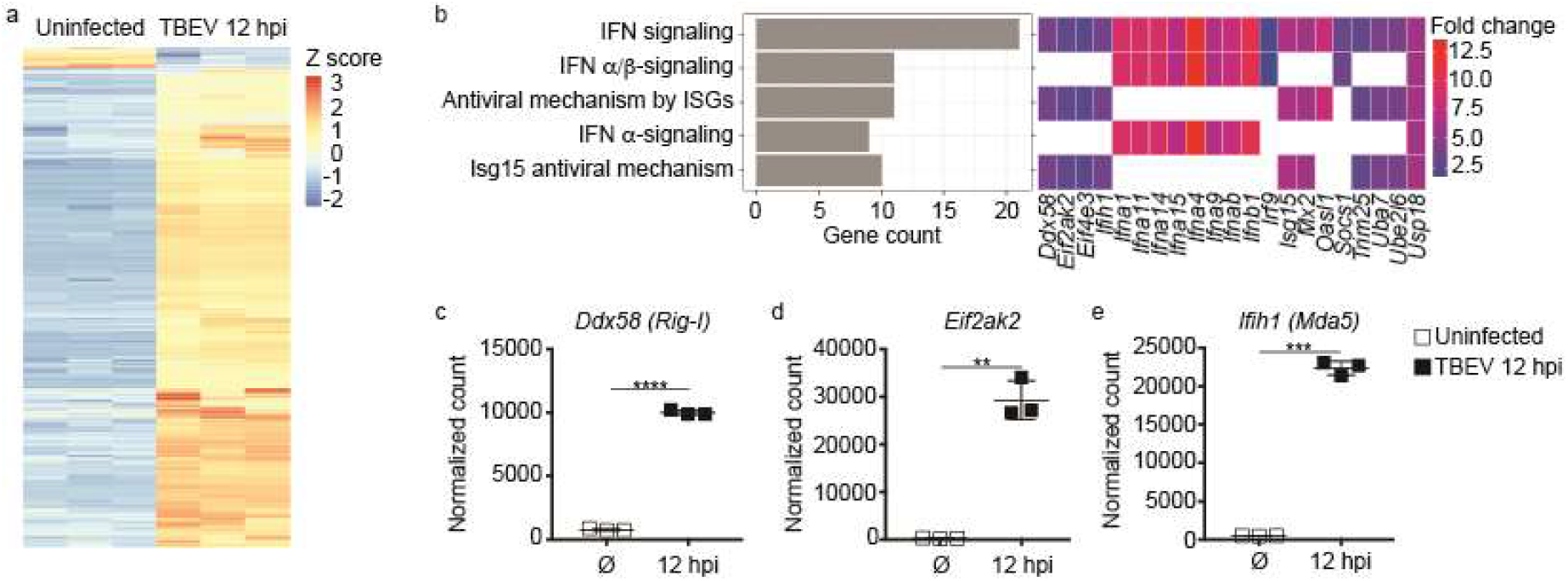
TBEV infected astrocytes show MAVS-dependent induction of early anti-viral responses. Primary murine astrocytes were either infected with TBEV Neudoerfl strain at MOI 1 or mock treated. At 12 hours post treatment, RNA-seq analysis was performed. **a**, Heatmaps showing relative expression of transcripts obtained from transcriptome analysis of mock treated (ø) or 12 h TBEV infected astrocytes (columns represent biological replicates). **b**, Enrichment analysis of significantly expressed genes in TBEV infected versus mock treated astrocytes, showing the top 5 enriched reactome pathways. On the x-axes, the gene counts associated with each pathway are shown. On the right side, the single genes associated with each pathway with the relative fold-induction in 12 h TBEV infected astrocytes compared to mock treatment. **c-e**, Graphs showing normalized reads of genes associated with IFN-related pathways in uninfected and 12 h TBEV infected astrocytes (each square represents a biological replicate, n=3; error bars indicate mean ± SD; **P< 0.01; ***P< 0.001; ****P< 0.0001; Two tailed Unpaired t test with Welch’s correction).

### MAVS-signaling is required to restrict TBEV replication in infected astrocytes

To further address the role of RLR-signaling, we analysed murine astrocytes isolated from mice that were deficient of MAVS, the adaptor of RLR-signaling. Upon exposure to TBEV strain Neudoerfl at MOI 1, approximately 30% of WT and more than 50% of MAVS ko astrocytes were infected, respectively, as indicated by the presence of TBEV replication foci (Fig. 3b). Quantification of the replication foci by determining their overall mean fluorescence intensity (MFI) revealed that 12 hpi WT astrocytes displayed low dsRNA signals, while MAVS ko astrocytes showed significantly higher signal intensities (Fig. 3c). These data suggested that MAVS ko astrocytes showed higher TBEV replication than WT astrocytes. Similarly, 24 hpi WT astrocytes were significantly less infected than MAVS ko astrocytes, with 50% of WT and 85% of MAVS ko astrocytes being infected, respectively (Fig. 3e). Moreover, quantification of TBEV replication foci 24 hpi revealed an overall higher virus replication than at 12 hpi, and MAVS ko astrocytes showed significantly higher numbers of TBEV replication foci than WT astrocytes (Fig. 3f). At 12 hpi, similar numbers of TBEV copies were detected in the SN of infected WT and MAVS ko astrocytes, whereas at 24 hpi the number of TBEV copies was significantly enhanced in the SN of MAVS ko astrocytes compared with the SN of WT astrocytes (Fig. 3g). These data indicated that TBEV replication was higher in MAVS ko astrocytes than in MAVS competent ones. Infected WT astrocytes produced high amounts of IFN-β already at 12 hpi, which did not further increase until 24 hpi (Fig. 3h). In contrast, MAVS ko astrocytes produced very little amounts of IFN-β at 12 hpi, whereas at 24 hpi IFN-β levels increased to moderate levels (Fig. 3h). These data supported the hypothesis that in TBEV infected astrocytes initially MAVS-signaling and later other mechanisms drive IFN-β expression.

**Figure 3:**
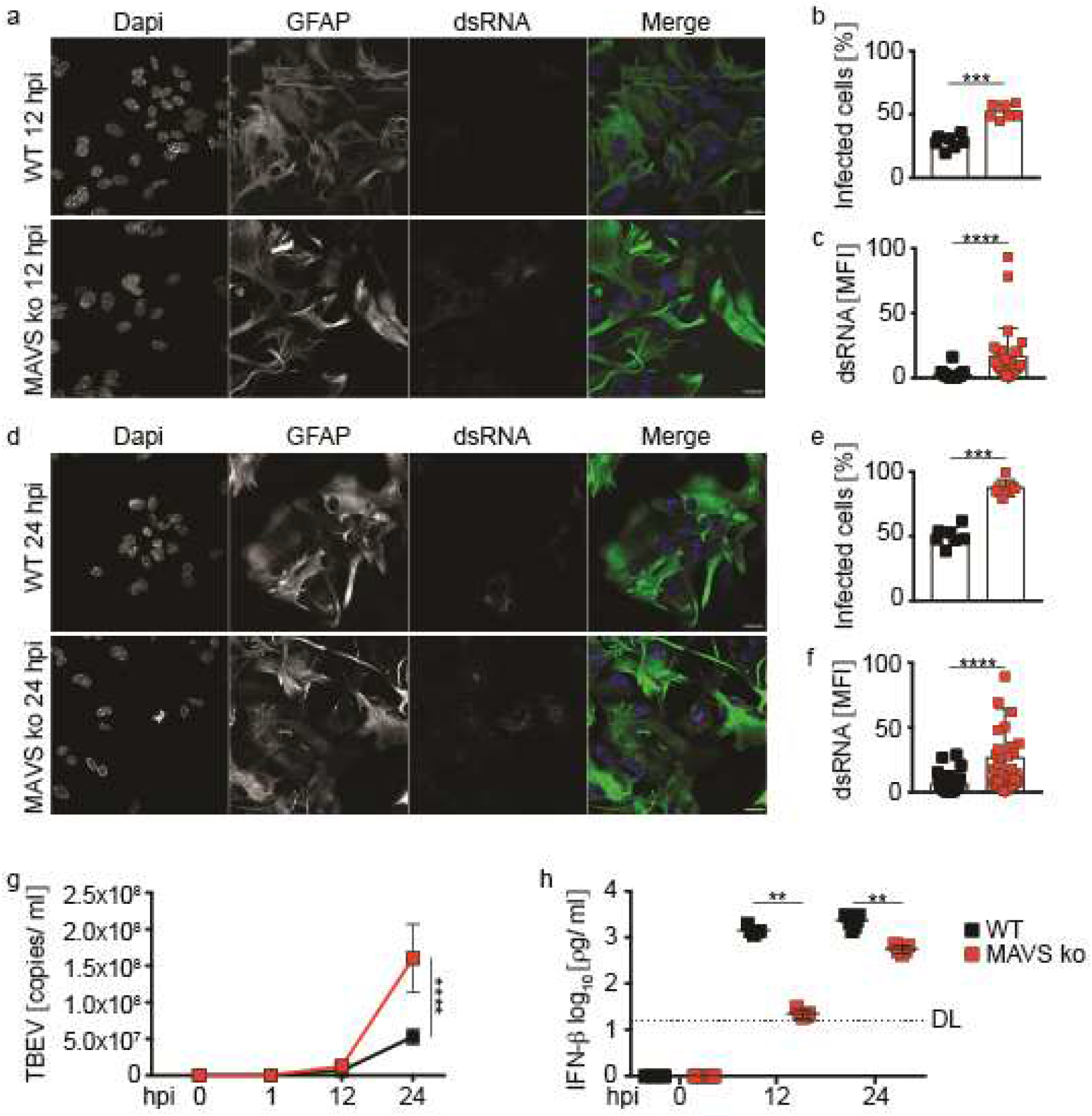
MAVS-deficient astrocytes fail to curtail TBEV replication. Primary murine astrocytes isolated from WT or MAVS ko mice were infected with TBEV Neudoerfl strain at MOI 1. **a, c**, At 12 and 24 hpi WT and MAVS ko astrocytes were immunolabelled with anti-GFAP, anti-dsRNA antibody and counterstained with DAPI and immunofluorescence microscopy was performed, scale bar 20 μm. **b, e**, Quantification of WT and MAVS ko TBEV infected astrocytes at 12 and 24 hpi, graphs displayed percentage of infected cells (n=6, N=3, combined data, bar plot indicates mean + SD). **c, f**, dsRNA mean fluorescence intensity (MFI) within infected cells as a measure of TBEV replication foci at 12 and 24 hpi (n=6, N=3, combined data, bar plot indicates mean + SD). **g**, TBEV copies per ml in the SN of WT and MAVS ko astrocytes at 0, 1, 12 and 24 hpi quantified by RT-qPCR (n≥ 5, combined data from 3 independent experiments). **h**, IFN-β protein expression in the SN of WT and MAVS ko astrocytes at 12 and 24 hours post TBEV infection determined by ELISA (n≥ 5, combined data from 3 independent experiments) DL= Detection limit (16 pg/ml), error bars indicate mean ± SD; *P< 0.05, **P< 0.01; Two-tailed Mann-Whitney test.

### Functional MAVS signaling by astrocytes is required for the induction of early IFN and ISG responses

Principle component analysis of transcriptome data of infected and uninfected WT and MAVS ko astrocytes showed that the transcriptomes of uninfected WT and MAVS ko astrocytes clustered closely together, whereas infection led to an increased variance at 12 hpi as well as 24 hpi (Fig. 4a). While uninfected controls displayed comparable gene counts independently of whether the analysed astrocytes were MAVS competent or deficient, 12 and 24 hpi WT astrocytes showed substantially higher gene counts than MAVS ko astrocytes (Supplementary Fig. 3a). TBEV infection induced upregulation of several genes in WT and MAVS ko astrocytes at 12 and 24 hpi (Supplementary Fig. 3b). Comparative analysis of differentially expressed genes (DEGs) at 12 hpi revealed 437 DEGs in WT astrocytes and only 212 DEGs in MAVS ko astrocytes. 206 transcripts were common between WT and MAVS ko infected astrocytes, while 267 DEGs were only found in infected WT astrocytes (Fig. 4b). Similarly, 24 hpi WT and MAVS ko astrocytes displayed 632 and 314 DEGs, respectively, of which 274 were common (Fig. 4b). To investigate the impact of MAVS-signaling on antiviral genes, a direct comparison of the transcriptomes of WT and MAVS ko infected astrocytes was performed. At 12 hpi, WT astrocytes showed higher induction of *Ifnb1, Ifnab, Ifna2-4-5* and ISGs than MAVS ko astrocytes (Fig. 4c). Similarly, at 24 hpi IFN genes and cytokines were found to be more abundantly induced in WT than in MAVS ko astrocytes (Fig. 4d). Furthermore, MAVS ko astrocytes showed decreased induction of various cytokines and chemokines at 12 and 24 hpi (Supplementary Fig. 3c). Interestingly, *Il1b* and *Cxcl5* were upregulated only in infected MAVS ko astrocytes, but not in WT astrocytes (Supplementary Fig. 3c). Since lower IFN-β protein levels were detected at 12 and 24 hpi in the SNs of MAVS ko astrocytes than in WT astrocytes, and at 24 hpi MAVS ko astrocytes produced similar amounts of IFN-β as WT astrocytes at 12 hpi (Fig. 3h), we next investigated whether MAVS ko astrocytes showed delayed induction of IFN genes. Indeed, 12 hpi MAVS ko astrocytes showed less abundantly induced *Ifnb1*, *Ifna2* and *Ifna4* genes than WT astrocytes (Fig. 4e), whereas the induction of IFN genes was stronger in MAVS ko astrocytes 24 hpi than 12 hpi (Fig. 4e). Similarly, at 12 hpi interferon stimulated genes such as *Isg15*, *Mx1* and *Rasd2* were less abundantly induced in MAVS ko astrocytes than in WT astrocytes, while at 24 hpi these genes were also induced in MAVS ko astrocytes (Fig. 4e). In line with these observations, the number of TBEV transcripts was significantly higher in MAVS ko astrocytes than in WT astrocytes at 24 hpi (Fig. 4f). Together, these data indicate that the lack of MAVS-signaling is associated with reduced induction of IFN-β, IFN-αs and ISGs, which allows enhanced viral replication that further impairs the intrinsic response of astrocytes to TBEV infection.

**Figure 4:**
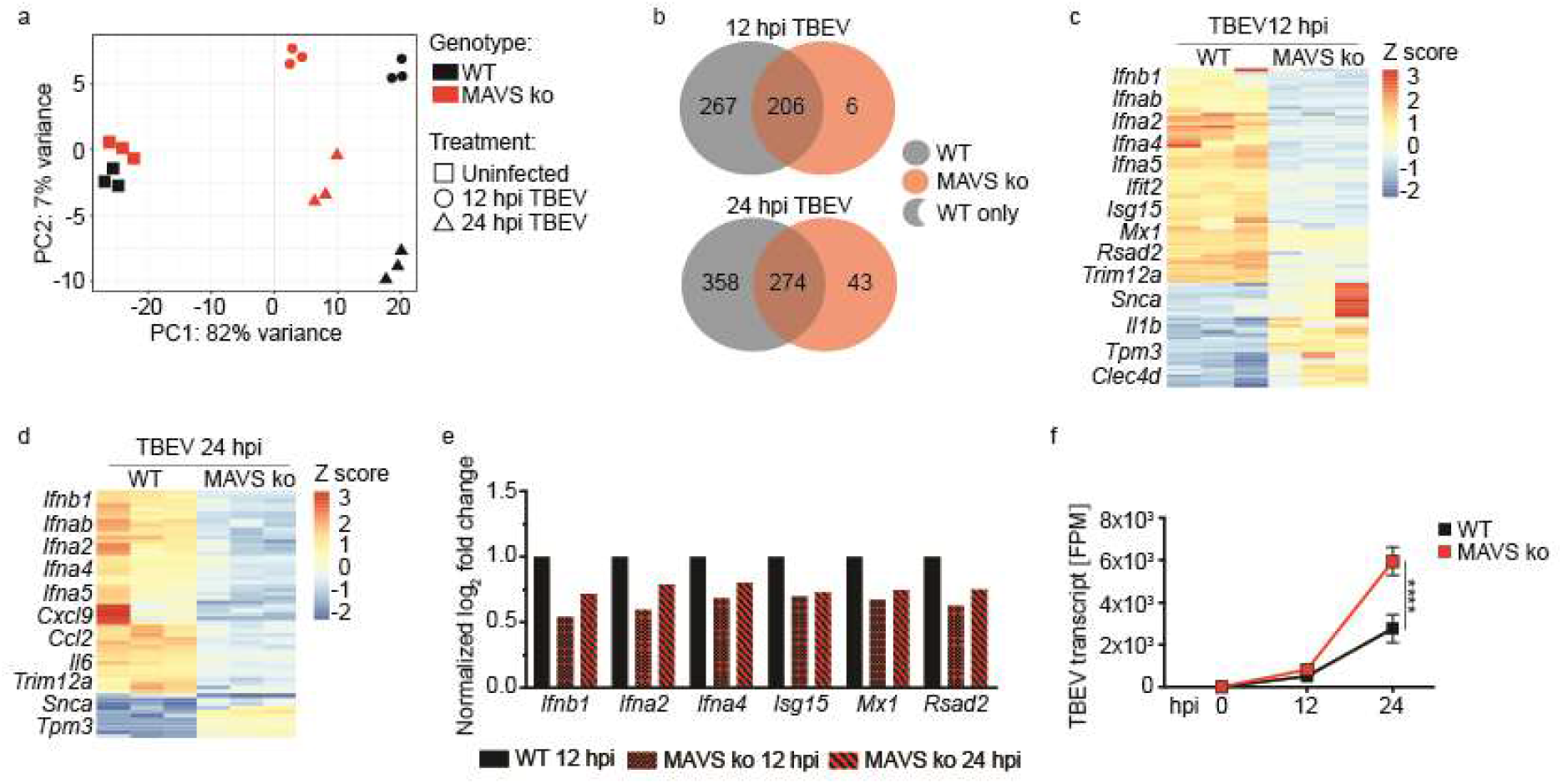
MAVS-deficient astrocytes show impaired anti-viral responses to TBEV infection. Primary murine astrocytes from WT or MAVS ko mice were either mock treated or infected with TBEV Neudoerfl strain at MOI 1. At 12 and 24 hours post treatment, cells were collected and RNA-seq analysis was performed. **a**, Principal component (PC) analysis of WT and MAVS ko astrocytes mock treated or TBEV infected for 12 and 24 hours (each dot represents one individual sample). **b**, Venn diagram showing the number of differentially expressed genes (DEGs) between the transcriptomes of WT and MAVS ko astrocytes at 12 and 24 hpi. **c**, Heatmaps showing relative expression of statistically significant transcripts obtained from transcriptome analysis of WT and MAVS ko astrocytes at 12 hpi (cutoffs: absolute log_2_ fold change > 2, each column represents transcripts obtained from one individual sample). **d**, Heatmaps showing relative expression of statistically significant transcripts obtained from transcriptome analysis of WT and MAVS ko astrocytes at 24 hpi (cutoffs: absolute log_2_ fold change > 2, each column represents transcripts obtained from one individual sample). **e**, Bar plot display *Ifnb1*, *Ifna2*, *Ifna4 Isg15*, *Mx1* and *Rasd2* gene expression in MAVS ko TBEV infected astrocytes at 12 and 24 hpi in comparison to TBEV infected WT 12 hpi. Log_2_ fold change values were normalized to TBEV infected WT 12 hpi with an assigned value of 1 (each column represents the mean value of normalized log_2_ fold change, n= 3, combined data). **f**, Graph depicts transcripts of TBEV in WT and MAVS ko astrocytes at 0, 12 and 24 hpi (n=3 combined data). TPM= transcript per million. ****P < 0.0001, 2way ANOVA Multiple comparisons test.

### IFN-β induction in infected astrocytes is dependent on sequential MAVS- and MyD88/TRIF-signaling

To further elucidate to which extent MAVS-signaling contributes to the induction of innate immune responses, pathways that were upregulated upon TBEV infection in WT, but not in MAVS ko astrocytes were investigated (Fig. 4a). This analysis revealed that at 12 hpi multiple pathways related to innate immunity were dependent on functional MAVS-signaling, including cytokine-mediated signaling, immune effector processes, regulation of innate immune responses, and the JAK-STAT cascade (Fig. 5a). At 24 hpi, MAVS-dependent signaling pathways were associated with adaptive immunity such as cell-cell adhesion, adaptive immune responses and lymphocyte differentiation (Fig. 5a). Interestingly, 24 hpi pathways related to innate immune responses were not dependent on MAVS-signaling, pointing towards the activation of MAVS-independent signaling. To investigate whether MyD88/TRIF-signaling contributed to the induction of protective IFN-β at later stages of *in vitro* TBEV infection, WT and MAVS astrocytes were treated with MyD88 and TRIF inhibitors, then infected with TBEV and IFN-β protein levels in the SNs were determined by ELISA. SNs of infected WT astrocytes that were not treated with inhibitors contained IFN-β at 12, 24 and 48 hpi (Fig. 5b), whereas SNs of infected MAVS ko astrocytes showed significantly lower IFN-β levels at 12 and 24 hpi than WT astrocytes and similar levels as WT astrocytes at 48 hpi (Fig. 5b). Upon treatment with MyD88 and TRIF inhibitors, infected WT astrocytes showed reduced IFN-β levels at 12 hpi that were further decreased at 24 hpi and undetectable at 48 hpi (Fig. 5c), whereas in the SNs of infected MAVS ko astrocytes no IFN-β was detectable at any of the tested time points (Fig. 5c). Since MAVS-independent pathways conferred the induction of late IFN-β, we hypothesized that TLR stimulation via MyD88/TRIF-signaling pathways might be relevant. Previous studies showed that astrocytes do not express the single stranded RNA sensor TLR7 (19). In contrast, *Tlr3* was upregulated upon TBEV infection of astrocytes, whereas this effect was stronger in WT than in MAVS ko astrocytes at 12 hpi (Fig. 5d). Furthermore, TRIF (*Ticam1*), the adaptor of *Tlr3,* was moderately upregulated in both infected WT and MAVS ko astrocytes (Fig. 5d). Similarly, MyD88 was upregulated in infected WT and MAVS ko astrocytes with a significantly higher induction in WT astrocytes at 12 hpi than in MAVS ko astrocytes (Fig. 5d). In conclusion, our data are compatible with the model of a bi-phasic induction of anti-viral responses in TBEV infected astrocytes, while at early time points MAVS-signaling and at later time points MyD88/TRIF-signaling is critically needed.

**Figure 5:**
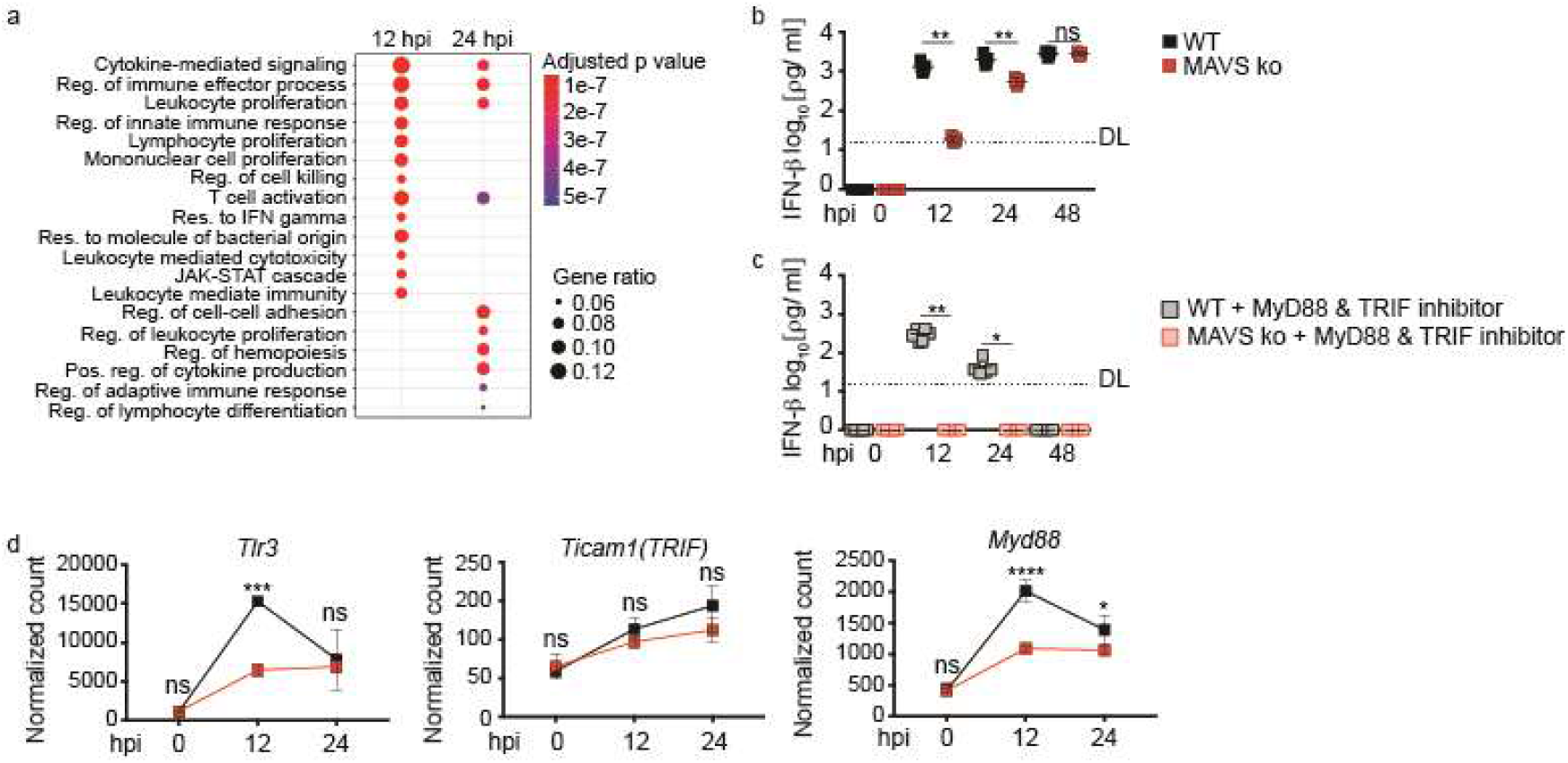
TBEV induced IFN-β responses of astrocytes initially depend on MAVS- and later on MyD88/TRIF-signaling. **a**, Pathway analysis showing top 20 downregulated GO terms imported from a gene set enrichment analysis (GSEA) of infected MAVS ko astrocytes compared with WT astrocytes at 12 and 24 hpi. The dot size indicates gene ratio, the number of genes within this GO term in comparison to the total number of upregulated genes. **b**, WT and MAVS ko astrocytes were infected with TBEV and IFN-β protein level were determined by ELISA at 0, 12, 24 and 48 hpi. **c**, WT and MAVS ko astrocytes were pre-treated with MyD88 and TRIF inhibitors (5 μM) for 8 h, then infected with MOI 1 of TBEV. At 0, 12, 24 and 48 hpi IFN-β protein level were determined by ELISA (Untreated sample are the same depicted in Fig.3h, for (**b**,**c**) n≥ 4, combined data; DL= Detection limit (16 pg/ml); error bars indicate mean ± SD; ns= not significant, *P< 0.05, **P< 0.01; Two-tailed Mann-Whitney test). **d**, Graphs showing *Tlr3, Ticam1 and Myd88* normalized gene counts in uninfected and 12 and 24 h TBEV infected WT and MAVS ko astrocytes (each square represents mean value of 3 biological replicate; error bars indicate mean ± SD; ns= not significant; *P< 0.05; **P< 0.01; ***P< 0.001; ****P< 0.0001; 2way ANOVA Multiple comparisons test).

## Discussion

Astrocytes are major IFN-β producers during CNS infection with a whole variety of RNA encoded viruses, including WNV (23), vesicular stomatitis virus (VSV (24)) and rabies virus (RABV (25)). Here, we observed that upon *in vitro* TBEV infection of murine brain-derived cell subsets, neurons showed the highest extent of infection and little IFN-β production, whereas astrocytes and microglia were less permissive to TBEV infection and produced significantly higher levels of IFN-β. Since astrocytes turned out to be the strongest IFN-β producers after TBEV infection, we further investigated the underlying mechanisms. Transcriptome analyses revealed that early after infection, signaling of MAVS, the adaptor of RLRs, is needed to induce IFNs and anti-viral responses in astrocytes. Consistently, astrocytes isolated from MAVS deficient mice showed increased TBEV replication as well as substantially decreased IFN-β induction early after infection. Interestingly, at later time points MAVS-independent and MyD88/TRIF-dependent mechanisms triggered almost normal anti-viral responses. These data support the hypothesis that TBEV infected murine astrocytes mount anti-viral responses in a bi-phasic manner that initially is driven by MAVS- and later by MyD88/TRIF-signaling.

In case of flavivirus infection*, in vivo* as well as *in vitro* experiments indicated that IFN responses are important to limit virus replication and to support recovery form the infection (26–29). Accordingly, TBEV infected murine astrocytes mount rapid IFN responses (23, 30), whereas IFNAR-deficient murine astrocytes showed increased virus replication and cytopathic effects (23). As a consequence of the induction of IFN responses and activation of JAK-STAT signaling, ISGs are induced (31, 32). Amongst ISGs that confer anti-viral effects during flavivirus infection, *Oas1* was shown to cleave viral mRNA (33). Similarly, viperin was reported to be induced in astrocytes, and to be involved in proteasome-dependent degradation of TBEV and Zika virus (ZIKV) NS3, thus restricting TBEV replication and spread (30, 34, 35). We found that TBEV showed sustained replication in MAVS ko astrocytes and that substantially reduced RNA levels of IFNs and ISGs, including *Oas1* and viperin, were induced. TRIM5a has previously been shown to contribute to the restriction of infection with TBEV and other flaviviruses in murine astrocytes (36). Interestingly, in our experiments *Trim12a*, which is the truncated murine homologue of human TRIM5a (37), was induced in TBEV infected WT astrocytes at early and late time points, whereas its induction was not detectable in MAVS ko astrocytes.

Earlier studies already suggested a key role of MAVS during LGTV infection, as indicated by MAVS ko mice that succumbed to systemic infection and showed higher viral loads and accelerated virus spread within the CNS (22). However, in these experiments, cytokine levels in brain homogenates of LGTV-infected MAVS ko mice were similar as in WT animals at 4 dpi (22), suggesting that by that time MAVS-independent mechanism must have been in place taking over cytokine induction within the CNS. Upon RNA virus infection, the viral RNA is recognized by TLRs or RLRs including RIG-I and MDA5 (38). RLR-signaling has previously been shown to play a key role in initiating immune responses upon Flavivirus infection such as WNV, DENV and others (39–41). Indeed, in case of *in vitro* WNV infection, RIG-I and MDA5 were shown to be engaged by viral PAMPs that arose during virus replication (42, 43). In line with these studies, we showed that induction of early anti-viral responses in astrocytes relies on MAVS-signaling, whereas at later stages of infection, also MyD88/TRIF-signaling was important. Amongst TLRs, TLR3 and TLR7 have previously been shown to be involved in Flavivirus sensing, which for downstream signaling deploy the adaptors TRIF and MyD88, respectively (44). However, from these TLRs astrocytes only expresses TLR3 (19). Whereas TLR3 signals via the adaptor molecule TRIF (45), MyD88 is the adaptor for all other TLRs and IL-1 receptor signaling (46, 47). While it is still unclear which viral PAMPs trigger TLR3 during Flavivirus infection, previous studies showed that in the case of WNV and DENV, TLR3 is involved in the induction of anti-viral responses (41, 48, 49). Also the inflammasome has been shown to contribute to the induction of antiviral-responses during WNV and DENV infection (50–52). However, the signal triggering IL-1β expression still has not been uncovered. Here we found that MAVS ko astrocytes show reduced and delayed responses to TBEV infection. Nonetheless, TBEV infected MAVS ko astrocytes upregulate *Tlr3, Ticam1* and *Myd88* genes as well as the gene encoding for IL-1β thus highlighting a possible explanation for the involvement of MyD88/TRIF signaling during the induction of late anti-viral responses. Further studies are needed to uncover whether late anti-viral responses of MAVS ko astrocytes involve the TLR3-TRIF and/or the IL-1β-MyD88 axis.

Similar to the mouse system, within TBEV infected human neurons, the virus replicates, spreads and induces cytopathic effects (53, 54). Human astrocytes show sustained TBEV infection and virus replication without the induction of cytopathic effects or cell death until 14 dpi (55, 56). Recently it was shown that human neurons and astrocytes differentially respond to TBEV infection. While neurons were more susceptible to the infection and showed reduced immune responsiveness, astrocytes were more resistant to infection and mounted stronger anti-viral responses. Moreover, astrocytes contributed to neuronal protection against TBEV infection (57). During TBEV infection of human astrocytes, the RLRs RIG-I and MDA5, but not TLR3, sense TBEV NS5 and activate IRF-3 signaling, which leads to the production of CCL5 (58). A recent study analyzing RNA expression profiles of TBEV infected human neuro-glia cultures, i.e., neuron and astrocyte cocultures, showed that in astrocytes the transcripts of *Ddx58, Oas2, IFN-*β*, Mx1, Trim5a* and *Rasd2* were even more abundantly induced than in neurons (57). We also observed upregulation of these transcripts in TBEV infected WT astrocytes, whereas in MAVS ko astrocytes the induction of the respective transcripts was significantly decreased.

Fares *et. al.* showed that at early stages of the infection human TBEV infected astrocytes show TLR3 expression that is close to basal levels, whereas at 24 hpi TLR3 expression is significantly enhanced (57). We also observed induction of TLR3 upon infection of both, WT and MAVS ko astrocytes. Interestingly, a previous study showed that in humans a functional TLR3 is a risk factor for the development of encephalitis upon TBEV infection (59). In contrast, mouse studies with other Flaviviruses such as WNV and ZIKV revealed a protective role of TLR-signaling (60, 61).

Taken together, upon *in vitro* TBEV infection of mouse-derived brain resident cell subsets, astrocytes are the main IFN-β producers. Early after infection, astrocytes are stimulated in a MAVS-dependent manner, presumably via RLR signaling, whereas at later time points MyD88/TRIF-signaling was needed. These observations highlight that over time single pathogens are not only sensed in different tissues by different cell subsets that deploy multiple sensing mechanisms (62, 63), but that even in one cell subset distinct sensing mechanisms may be relevant at different times after infection. It will be a matter of future research to address whether biphasic virus sensing during TBEV infection similarly applies to human astrocytes.

## Acknowledgements

We thank Jennifer Skerra and Kira Baumann for help with mouse breeding and screening. We thank Carla Schmutte for excellent technical support. This study was supported by funding from the Helmholtz Association (Zukunftsthema “Immunology & Inflammation” (ZT-0027); https://www.helmholtz.de/en/), by the Niedersachsen-Research Network on Neuroinfectiology (N-RENNT) of the Ministry of Science and Culture of Lower Saxony, Germany, and by the Federal Ministry of Education and Research (BMBF) under project number 01KI1719 as part of the Research Network Zoonotic Infectious Diseases (V.B. and I.S.).

## Author contribution

Conceptualization, L.G., I.S. and U.K.; Methodology, L.G., V.B., A.P., F.M., O.G., C.K.P., I.S., and U.K.; Investigation, L.G., V.B., A.P., F.M., O.G., I.S.; Data Discussion: L.G., V.B., F.M., A.P., O.G., V.D., M.K., K.J., M.S., I.S. and U.K.; Writing – Original Draft: L.G., I.S. and U.K., Supervision: I.S. and U.K.; Funding Acquisition: M.S., I.S. and U.K.

## Material and method

### Virus propagation and quantification

TBEV strain *Neudoerfl* was provided by Dr. Gerhard Dobler and propagated in A549 cells. The cells were infected with TBEV at MOI 0.01 using high-glucose Dulbecco’s modified Eagle’s medium (DMEM) supplemented with penicillin (10,000 U/mL), streptomycin (10 mg/mL) and L-glutamine (200 mM) (Sigma). After one hour of incubation, the inoculum was removed and exchanged by DMEM supplemented with 2% fetal calf serum (FCS), penicillin and streptomycin, and L-glutamine. The supernatant was collected after five days and virus titration was performed on A549 cells. Viral RNA was extracted from culture supernatants with Qiagen Viral RNA Mini kit and quantification of genome copies was performed by probe-based quantitative RT-PCR according to Schwaiger and Cassinotti (64).

### Mice

C57BL/6 (Wild type or WT, Envigo) and B6.STOCK-Mavs(tm1Tsc) (Mavs^−/−^, MAVS KO) (65) mice were bred under specific pathogen-free conditions in the central mouse facility of the Helmholtz Centre for Infection Research, Brunswick, and at TWINCORE, Centre for Experimental and Clinical Infection Research, Hanover, Germany. Mouse experimental work was carried out in compliance with regulations of the German animal welfare law.

### Murine primary CNS resident cell isolation and culturing

Neurons were isolated and cultivated from mouse embryos at embryonic day 12.5, cortex free of meninges was dissected on ice cold PBS using a stereomicroscope (Nikon, SMZ45T), The cortex was collected in ice-cold Hanks Balanced Salt Solution (HBSS, Gibco) and minced with fine dissection scissors. Next, enzymatic digestion was performed, the minced tissue was incubated in 500 μL of trypsin-EDTA (Biochrome) and 2 μl of DNase (Roche) for 8 minutes at 37 °C. 4 mL of DMEM medium (Gibco) was then added and mechanical dissociation with 1 mL pipet was performed. The solution containing the dissociated cortex was then resuspended in 2 mL of neurobasal complete medium (Gibco), containing media supplements (1x B-27, Gibco; 1x N2, Gibco; 1% Pen/Strep, Capricorn; 1% GlutaMax, Gibco) and neurons were seeded in poly-L-lysine (Sigma) coated 24-well plates or on poly-L-lysine coated glass coverslips at a density of 2.5 × 10^5^ cells per well. Neurons were kept in culture for 7 days before infection and fresh neurobasal complete media was added every second day.

Microglia and astrocytes were isolated as previously described (66); in brief, brains from newborn mice were dissected at day 3 post birth, meninges and blood vessels were removed using a stereomicroscope (Nikon, SMZ45T). Next, the brains were minced and digested with 0.1% trypsin (Sigma-Aldrich) and 0.25% DNAse (Roche), single cell suspensions were seeded in precoated poly-L-lysine flasks in DMEM with 1% penicillin / streptomycin (Gibco) supplemented with 10% FCS. Medium was changed on day 1 and subsequently every 3 days. Microglia were collected after 9-11 days of culturing by shaking the flasks at 37 °C (180 rpm) in an orbital shaker for 40 minutes and plated the microglia at a density of 2.5 × 10^5^ cells per well into a 24-well plate or on glass coverslips. Microglia was used the following day for experiments. Astrocytes were isolated after removing remaining microglia and oligodendrocyte by over-night shaking at 37 °C (170 rpm), followed by cytosine arabinoside (AraC, Sigma-Aldrich) treatment for 3 days. Astrocytes were detached by trypsin-EDTA (bio-sell) treatment and plated into a 24-well plate or on glass coverslips at a density of 2.5 × 10^5^ cells per well. Medium was changed 24 hours after seeding and then used for infection.

### Infection and treatment of CNS primary cells

All cell types were inoculated with a MOI of 1 of TBEV in serum- and antibiotic free DMEM medium for 1 hour at 37 °C, then washed with warm PSB and incubated with the respective culture media. MyD88 and TRIF inhibition experiments was performed by pre-treating astrocytes with 5 μM of Pepinh-MYD and Pepinh-TRIF (invivogen) for 8 hours. At 12 and 24 hours post infection, cell-free SN was collected and stored at - 80 °C for IFN-β analysis and TBEV copy number determination. The adhering cells for RNA analyses were then harvested and lysed by TRIzol (Thermo Fischer) and stored at −80 °C.

### Immunocytochemistry and microscopy

Coverslips were fixed in 4% paraformaldehyde for 20 minutes at RT, than washed and stored in PBS. Non-specific binding of antibodies was inhibited by incubation for 1 hour at room temperature with blocking buffer containing 3% donkey serum (Sigma-Aldrich) and 0.3% Triton (Sigma-Adrich) in PBS. Next, the primary antibodies in blocking buffer were added for 2 hours at 4 °C. Excess antibodies were removed through 3 washing steps with PBS before incubation for 1 hour at room temperature with the secondary antibodies in blocking buffer. After washing, nuclei were counterstained with DAPI (4’,6-diamino-2-phenylindole, 1:10.000, Invitrogen) and the coverslips were mounted with mounting medium (DAKO) on glass slides. Pictures were taken on a Leica microscope SP8 using LAS-X software and analyzed by Fiji. Primary antibodies: mouse anti-mouse β-III tubulin (Tuj1) (Biolegend; 1:400), mouse anti-mouse GFAP (Millipore; 1:400), goat anti-mouse Iba-1 (Abcam; 1:200) and dsRNA (J2, Scicons English and Scientific Consulting, TBEV replication foci; 1:500). Secondary antibodies: donkey anti-rabbit AF568 (Invitrogen; 1:500), donkey anti-mouse AF568 (Invitrogen; 1:500), donkey anti-goat AF647 (Invitrogen; 1:500) and donkey anti-mouse AF 647 (Invitrogen; 1:500).

### IFN-β analysis

IFN-β protein levels were determined from cell-free SNs of control and TBEV infected cells, using the VeriKine Mouse Interferon ELISA kits (PBL Assay Science) and VeriKine Mouse Interferon ELISA High Sensitivity kits (PBL Assay Science) accordingly to the manufacturer’s handbook.

### RNA isolation and sequencing

RNA was isolated from WT and MAVS ko astrocytes either untreated or TBEV infected for 12 or 24 hours. RNA was extracted using Direct-zol RNA Miniprep Plus Kit (Zymo Research) according to the manufacturer’s instructions. RNA quality and integrity was assessed with Bioanalyzer 2100 (Agilent) and samples with RIN value > 8.0 were selected for further processing. Sequencing libraries were constructed and processed on Illumina NovaSeq-600 platform with 50 bp paired-end reads configuration. Quality control of the sequenced raw FASTQ-files was performed with FastQC software (version 0.11.9) and mapped to Ensembl mouse genome reference version GRCm38 (mm10) using STAR (67) (version 2.5.4b). Gene abundance was also determined by STAR, and differential expression analysis was performed with the DESeq2 package (68) in the R environment, setting the FDR to 0.05. Functional annotation and enrichment analysis was performed, as previously described (69), on the sets of differentially regulated genes using the Gene Ontology and Reactome databases.

## Supplementary information

**Figure S1:**
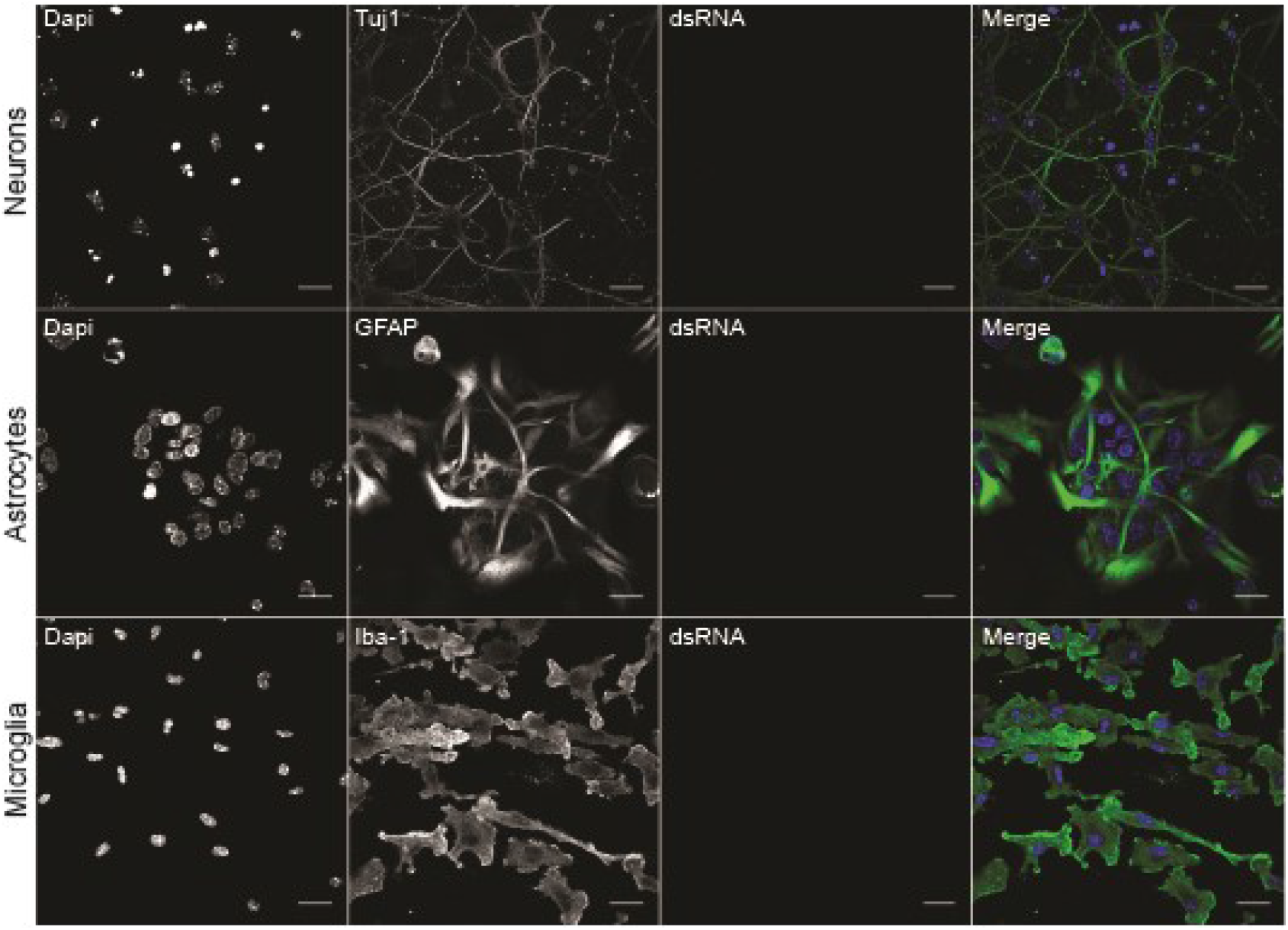
Morphological analysis of untreated murine primary CNS resident cells. Primary murine neurons, astrocytes and microglia from C57BL/6 (WT) mice were mock-treated and at 12 h post treatment, cells were immunolabelled with either anti-Tuj1 (neurons), anti-GFAP (astrocytes), or anti-Iba1 (microglia) and dsRNA (J2, Scicons English and Scientific Consulting, TBEV replication foci) and counterstained with DAPI. Scale bar 20 μm.

**Figure S2:**
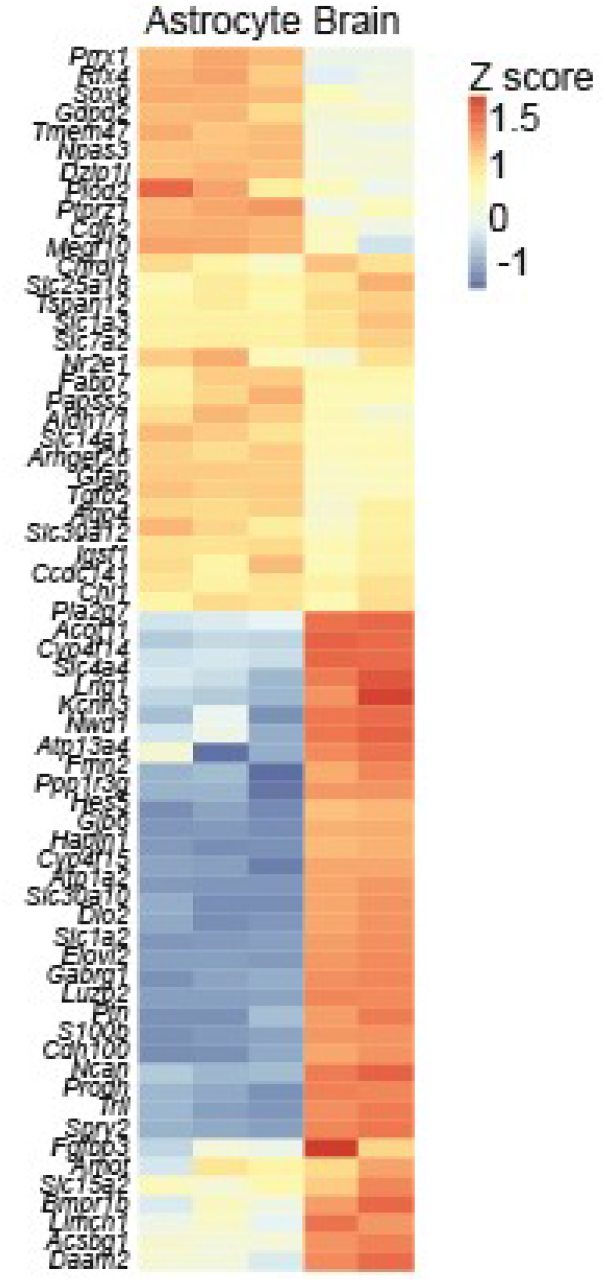
Verification of astrocyte identity by evaluation of astrocyte-specific transcripts. Relative expression of transcripts obtained from uninfected WT astrocytes compared with transcriptome from the total CNS, highlighting specific transcripts for astrocytes based on a previously published database (70), (each column represents an individual sample).

**Figure S3:**
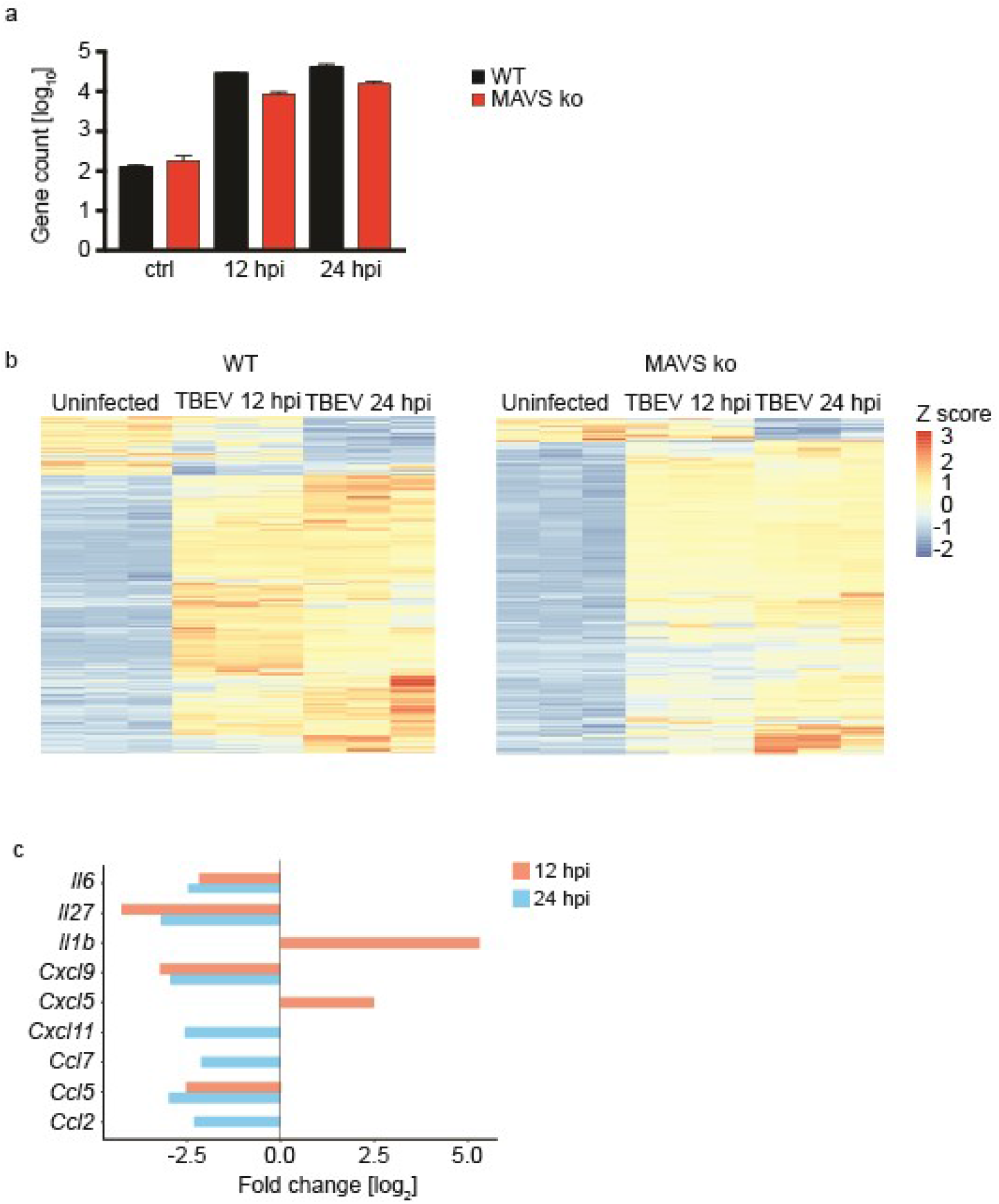
Upon TBEV infection, MAVS-deficient astrocytes show lower induction of transcripts than WT astrocytes. **a**, Comparison of total gene counts in uninfected (ctrl) and TBEV infected WT and MAVS ko astrocytes at 12 and 24 hpi. **b**, Heatmaps showing relative expression of transcripts obtained from transcriptome analysis of WT and MAVS ko astrocytes either uninfected or infected for 12 and 24 hours with TBEV (each column represents an individual sample). **c**, Log_2_ fold change of significantly altered cytokine and chemokine transcripts of infected MAVS ko astrocytes relative to WT astrocytes. Cutoffs: absolute log_2_ fold change>2.

